## Supplementary Figures File for "Assembly of the infant gut microbiome and resistome are linked to bacterial strains in mother’s milk": Supplementary_information.docx

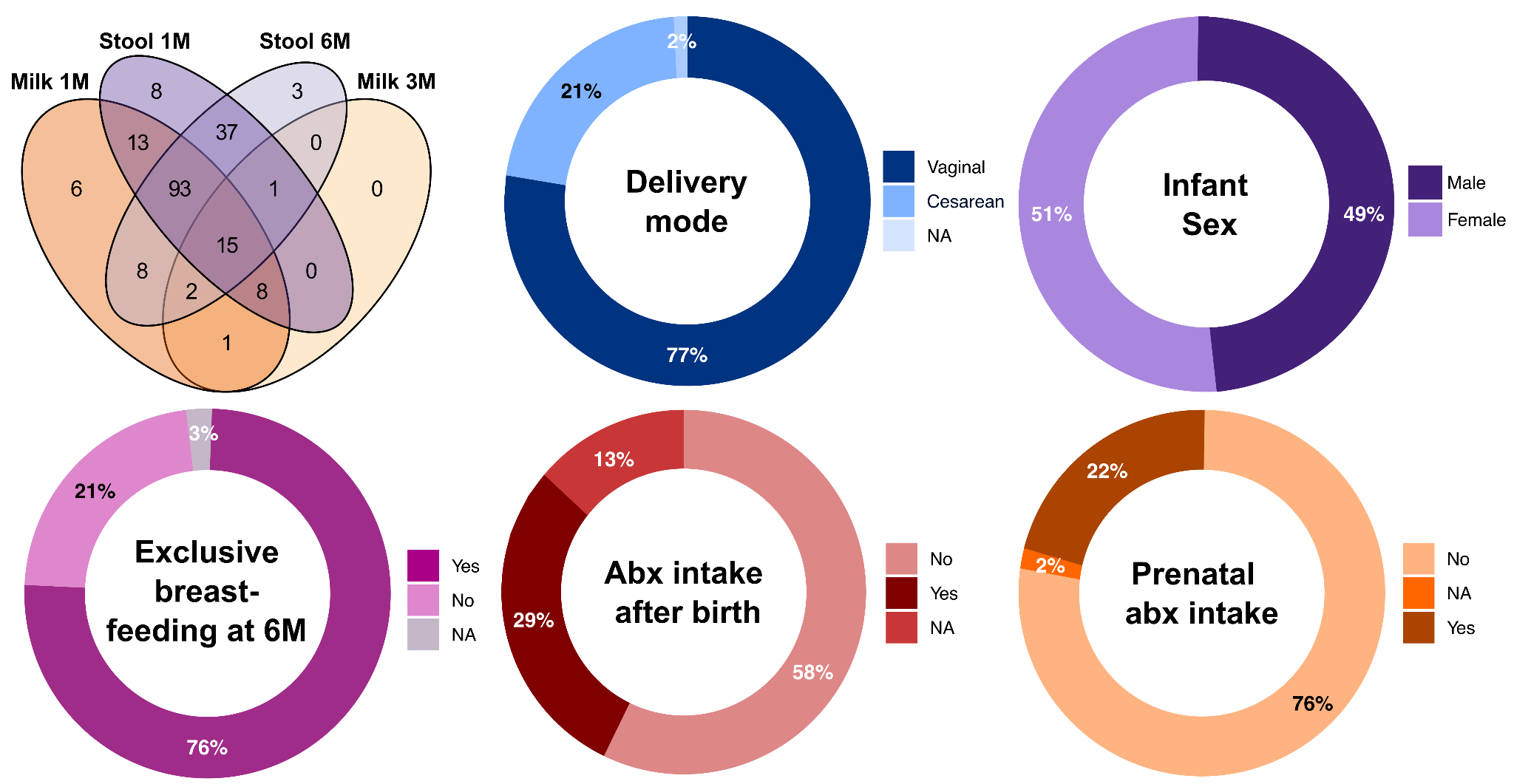

**Extended Data 1.** **Overview of the study metadata.** Number of mother infant pairs for which samples were collected across body sites and collection timepoints, singularly taken or in combinations. Relevant metadata available for the MILk cohort, including delivery mode, infant sex, exclusive breastfeeding at 6 months of age, and antibiotic (abx) intake pre- and after-partum. All infants were exclusively breastfed at 1 month of age.

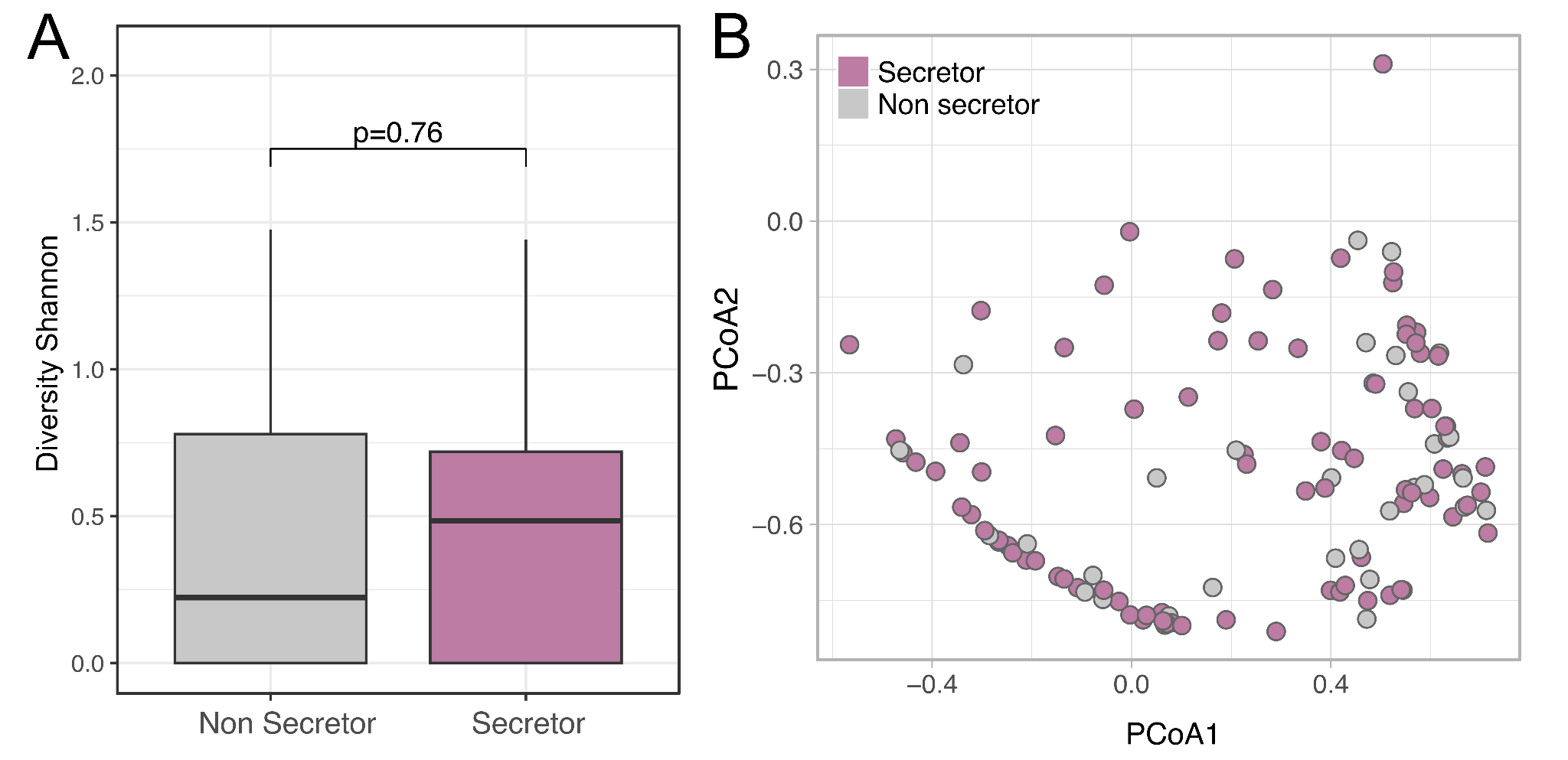

**Extended Data 2.** **Milk microbiome diversity and composition by secretor status**. (A) Alpha and (B) beta diversity of milk microbiome samples collected at 1 month postpartum stratified by maternal secretor status. See Methods for definition of secretor vs non secretor status. P-value calculated using paired t-test. Ordination plot based on Bray-Curtis distance between samples, colored by maternal secretor status. Milk samples collected at 3 months were not included.

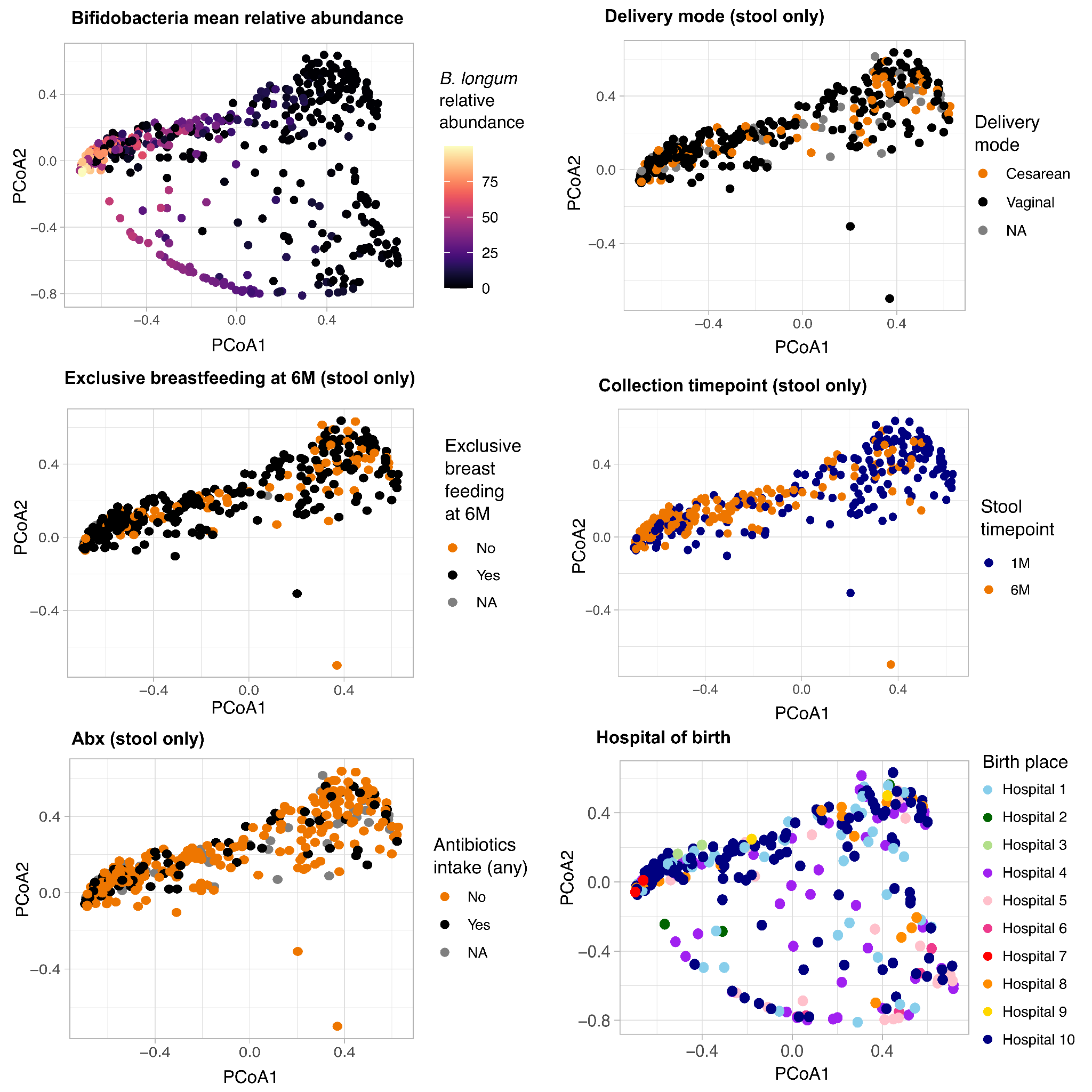

**Extended Data 3.** **Ordination plot coloured by relevant metadata.** All body sites and collection time points are included unless otherwise specified. PCoA of hospitals includes only the samples for which the birth hospital or clinic name is known.

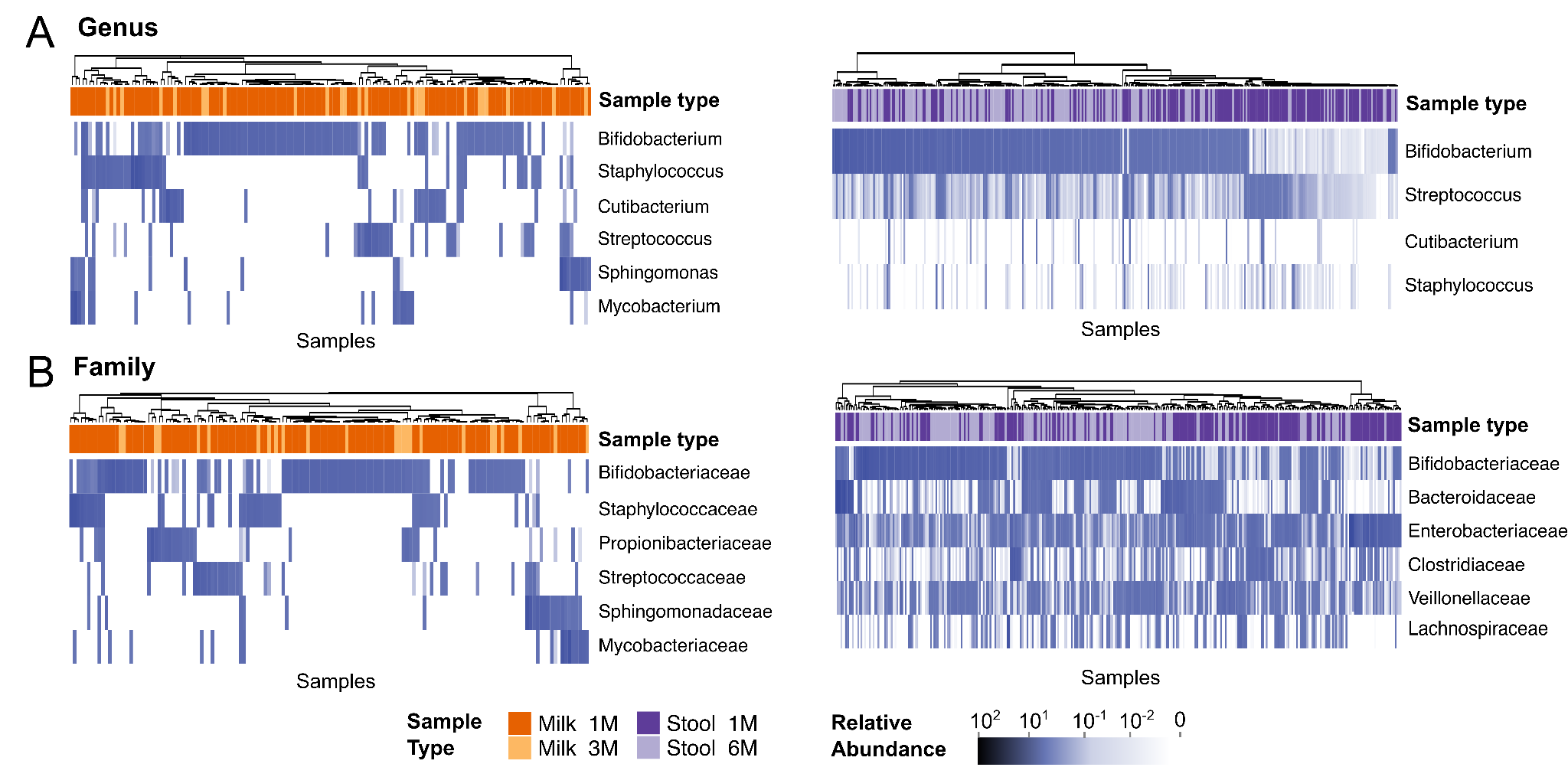

**Extended Data 4.** **Milk and infant stool microbiome composition**. Most prevalent bacterial genera (A) and families (B) in milk (left) and infant stool samples (right).

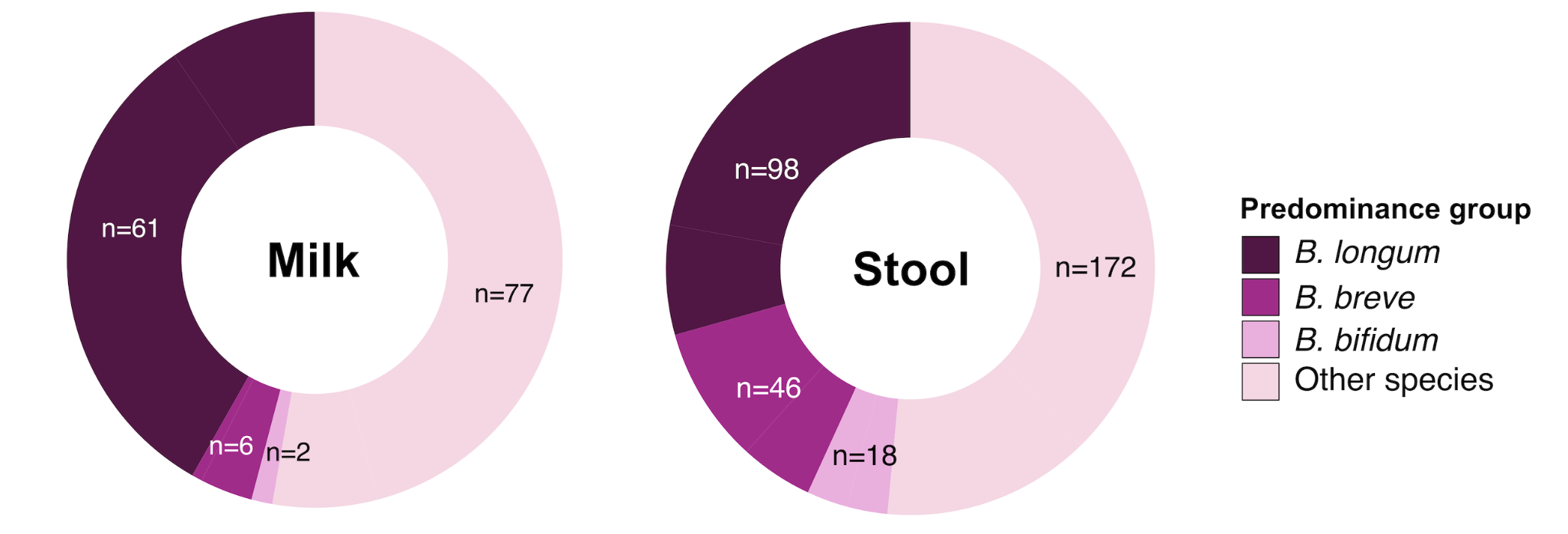

**Extended Data 5.** **Predominance groups in milk and infant stool samples.** Samples distributions across the four predominance groups and their associated metadata.

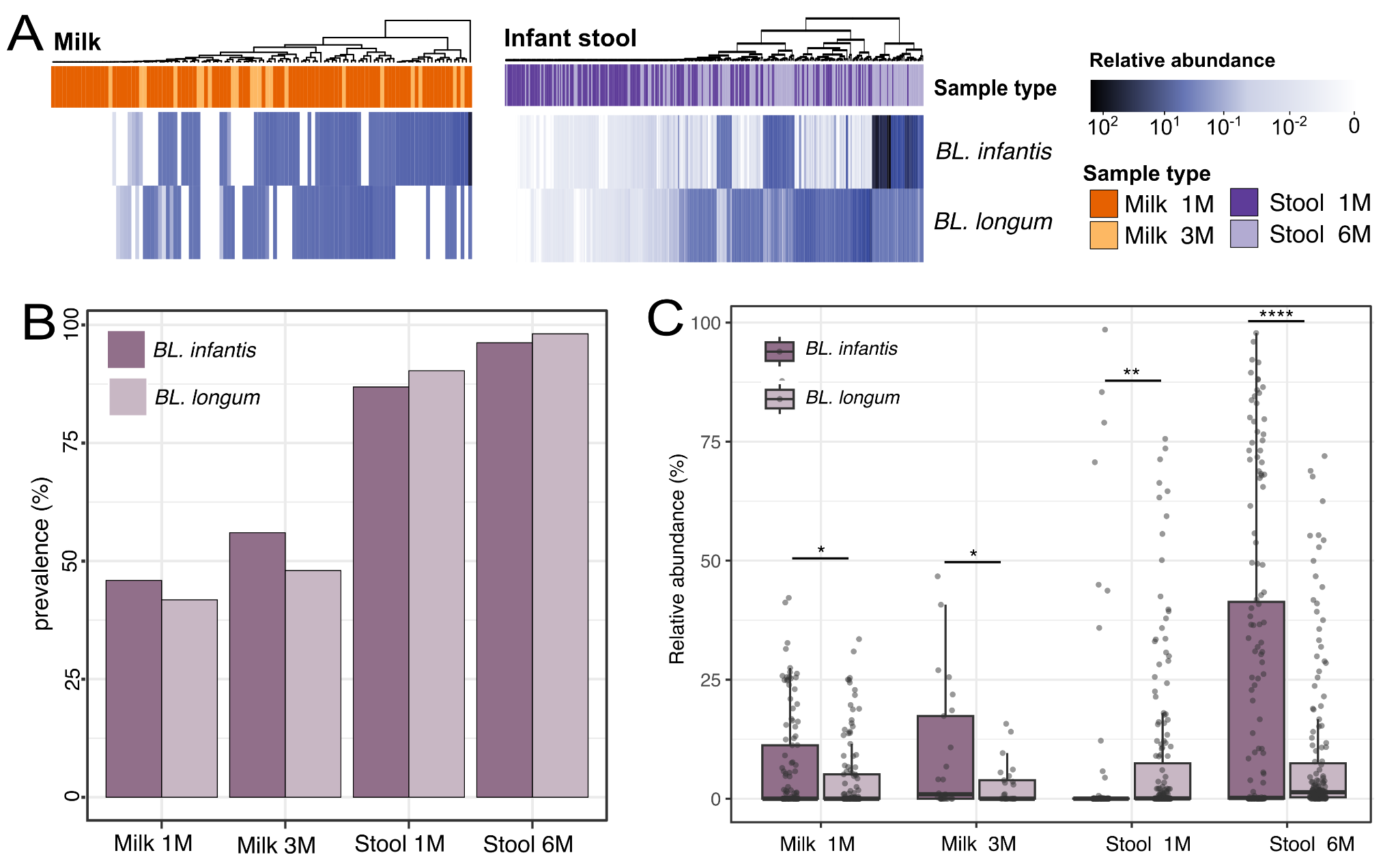

**Extended Data 6.** **Prevalence and abundance of *Bifidobacterium longum subspecies***. (A) Overview of *B. longum* subspecies in milk and infant stool samples, (B) their prevalence and (C) their relative abundance across sample types and over time. **** for p ≤ 0.0001, ** for p ≤ 0.01 and * for p ≤ 0.05. P-values calculated with paired t-test and adjusted with Bonferroni correction. *BL. infantis* = *B. longum* subsp. *infantis*; *BL. longum* = *B. longum* subsp. *longum*.

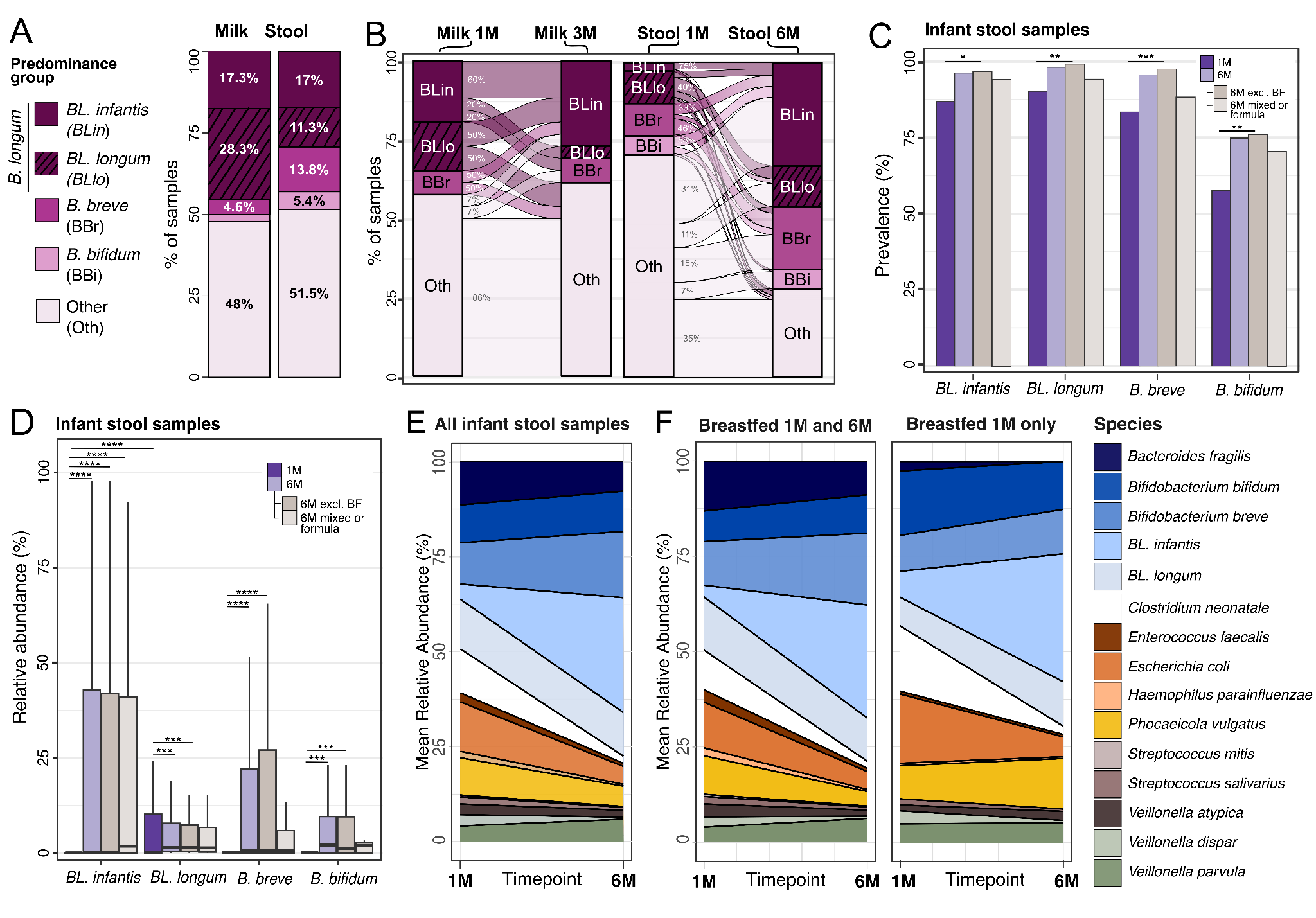

**Extended Data 7.** Predominance groups and *B. longum* subspecies overview. (A) Prevalence of each predominance group in milk and infant stool samples and (B) the transition of samples between predominance groups over time. Each sample is assigned one of four predominance groups indicated by the different colors. *B. longum* group is further split into its two most prevalent subspecies (*BL. infantis* and *BL. longum*). (C) Prevalence of *Bifidobacterium* species and subspecies at one and six months of age. Prevalence at six months is further stratified by exclusive breastfeeding status (n=121 for exclusive breastfeeding; n=34 for mixed/formula diet). Samples lacking breastfeeding information at six months were excluded from the stratified subgroups (n=4). Bonferroni adjusted p-values were calculated using Fisher’s exact test. (D) Relative abundance of bifidobacteria in infants at one and six months of age. Relative abundances at six months of age are further stratified by exclusive breastfeeding status. Relative abundances were calculated only on samples in which the species was present. P-values calculated using Wilcoxon rank sum, * for p ≤ 0.05, ** p ≤ 0.01, *** for p ≤ 0.001 and **** for p ≤ 0.0001. Reported p-values are adjusted using Bonferroni method. (E) Species persistence in the infant gut across all samples and (F) stratified by breastfeeding at six months. *BL. infantis* = *B. longum* subsp*. infantis*; *BL. longum* = *B. longum* subsp*. longum*.

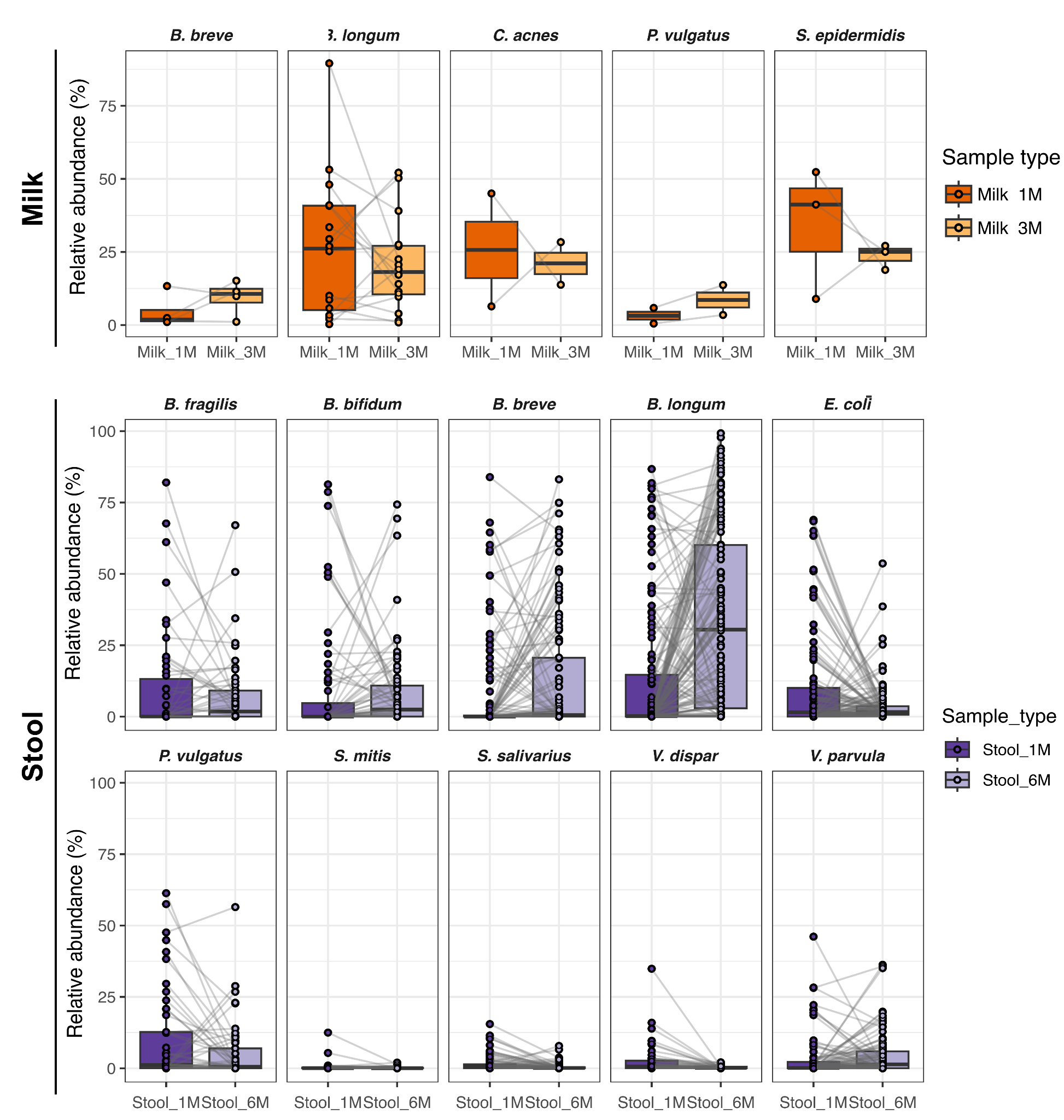

**Extended Data 8.** **Longitudinal trajectories of species-specific relative abundances.** Individual species-specific relative abundance trajectories over time, for milk and infant stool samples. Only mothers (for milk samples) or infants (for stool samples) for which both timepoints were collected are included.

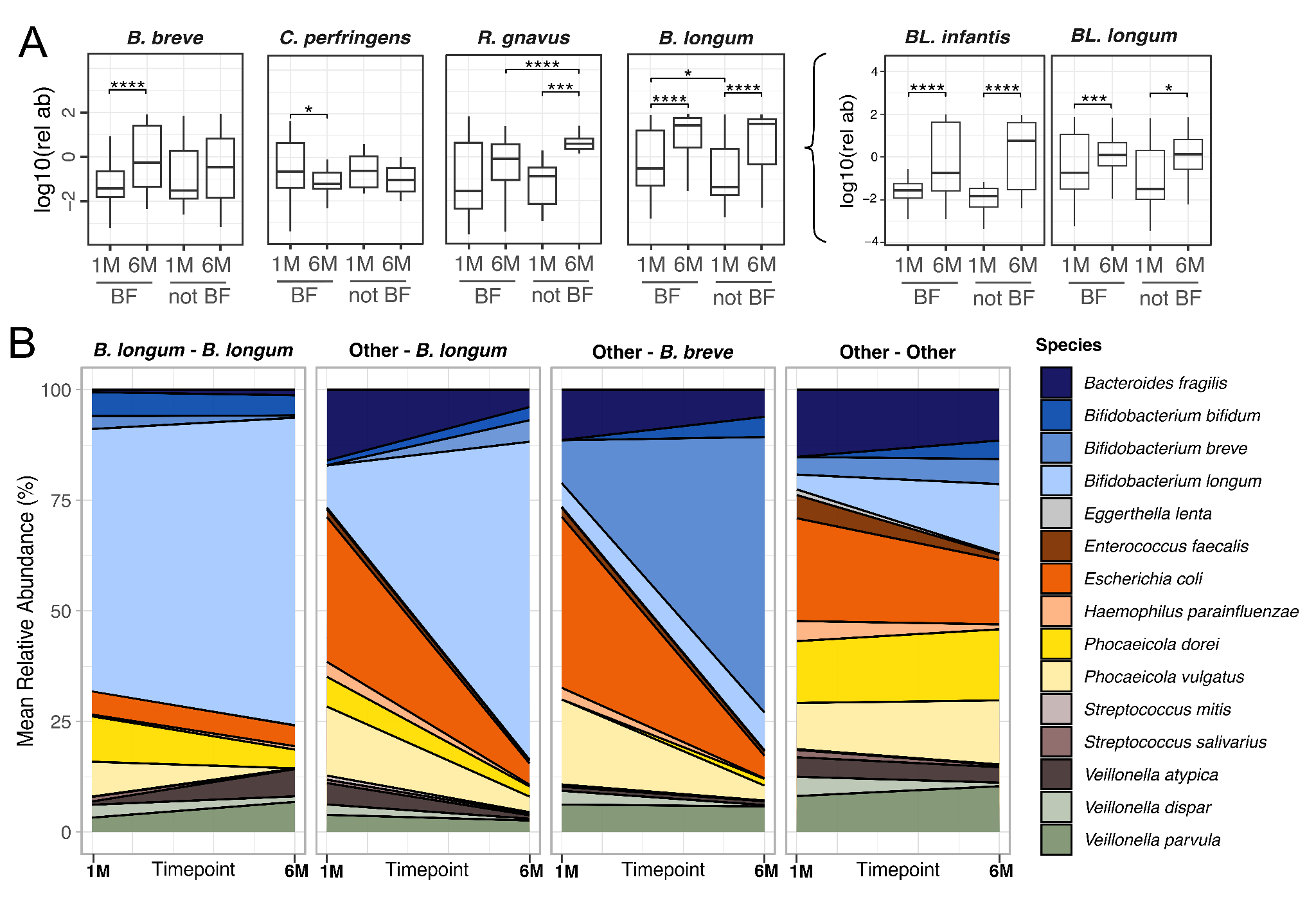

**Extended Data 9.** Differential abundance and persistence of microbial taxa in the infant gut microbiome. (A) Relative abundance of some of the differentially abundant species between one and six months when divided by breastfeeding (BF) at six months. *BL. infantis* = *B. longum* subsp. *infantis*; *BL. longum* = *B. longum* subsp. *longum*. **** for p ≤ 0.0001, *** for p ≤ 0.001, and * for p ≤ 0.05. P-values calculated with paired t-test and adjusted with BH correction. (B) Species persistence in the infant gut microbiome over time, stratified by the type of transition between predominance groups from one to six. Only samples with both time points available and transition types with more than ten samples per type were included.

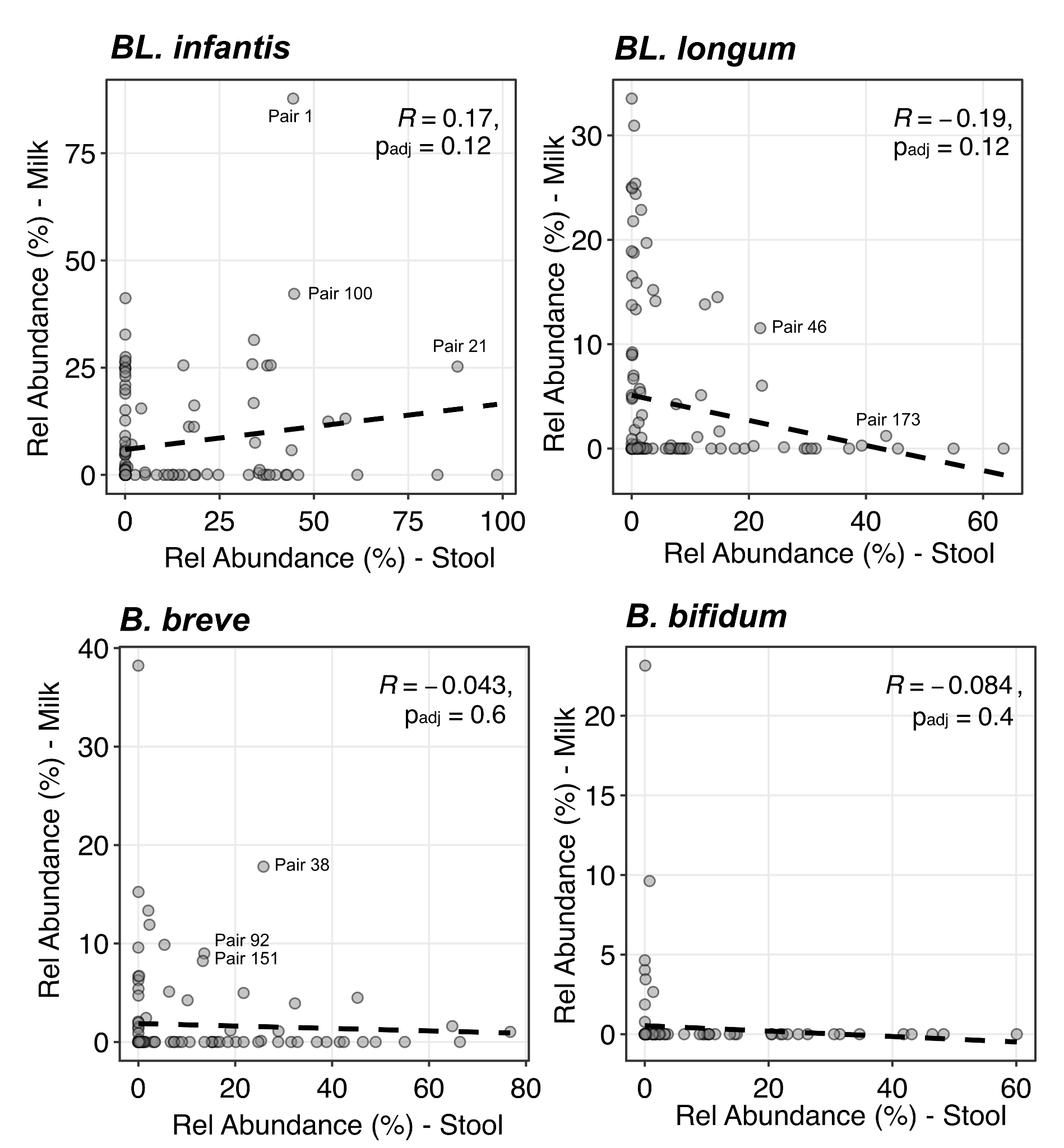

**Extended Data 10**. **Correlation between the relative abundance of specific bifidobacteria in milk and infant stool samples.** Each dot is a mother-infant pair. Mean relative abundances were calculated for infant stool samples at one and six months of age (x axis) and milk samples at one and three months postpartum (y axis). Only mother-infant pairs for which at least one milk and one infant stool sample were collected and passed preprocessing were included. R coefficients and p-values are reported. Correlation was calculated using Pearson. Bonferroni-adjusted P-values are reported.

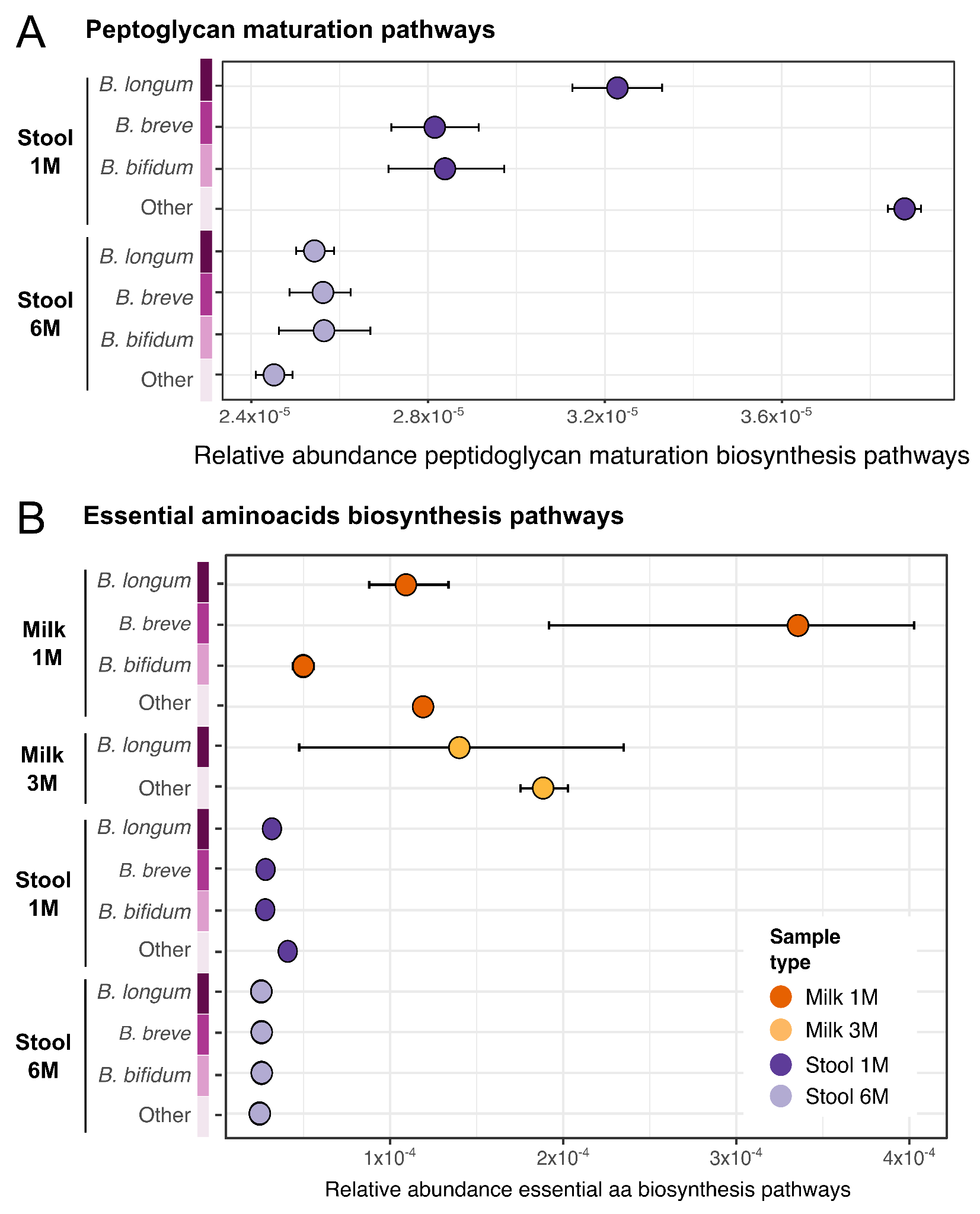

**Extended Data 11**. **Relative abundance of specific functional pathways across sample types and stool predominance groups**. Relative abundance of pathways associated with (A) peptidoglycan maturation and (B) essential amino acids biosynthesis across sample types and (stool) predominance groups. CI at 95%, bootstrapping n=1000.

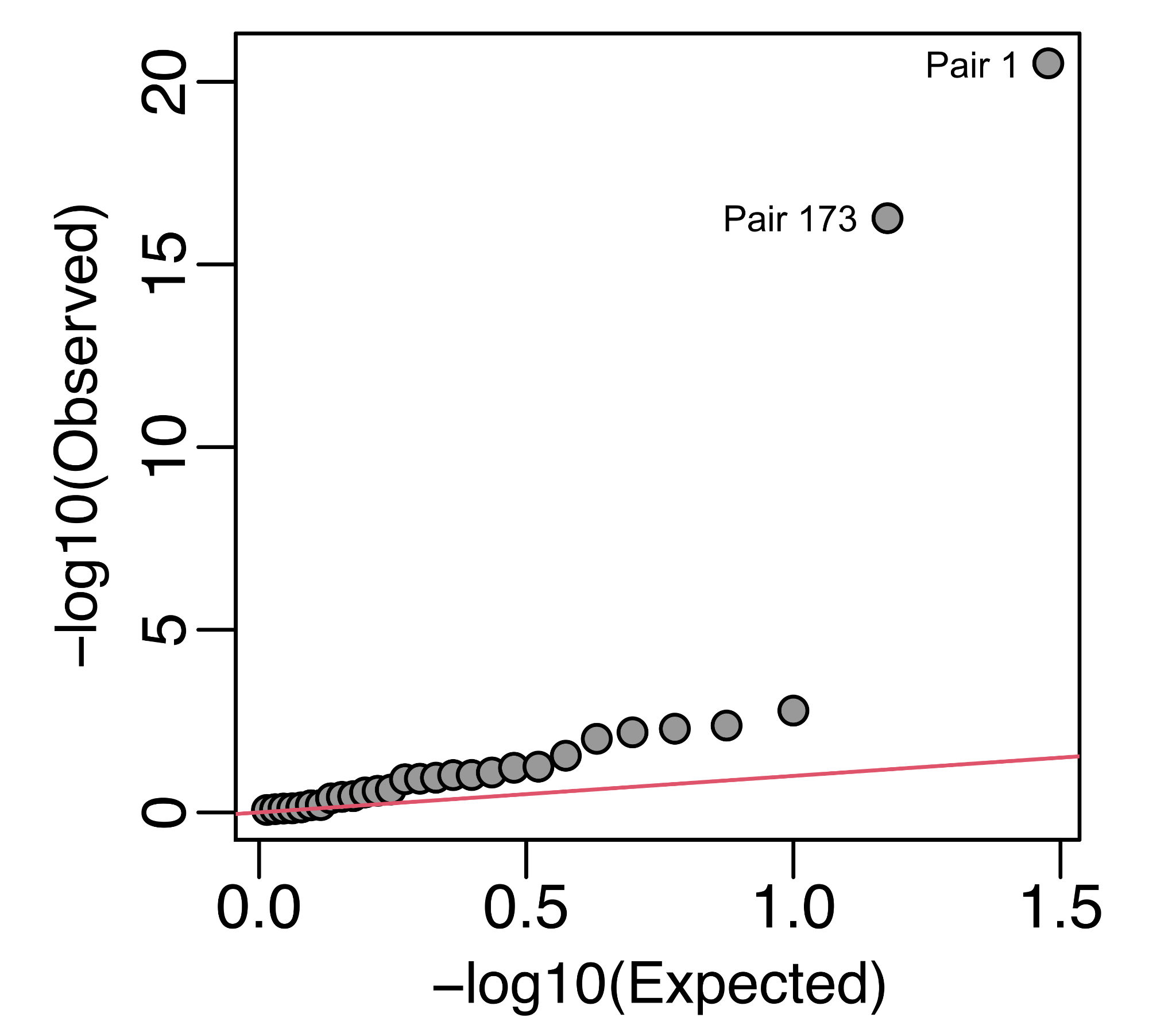

**Extended Data 12**. **Q-Q plot of Spearman’s correlation p-values for shared metabolic pathways between milk and infant stool across mother-infant pairs.** Each dot represents the p-value for one mother-infant pair. Mother-infant pairs 1 and 173 are highlighted. The red line indicates a uniform distribution.

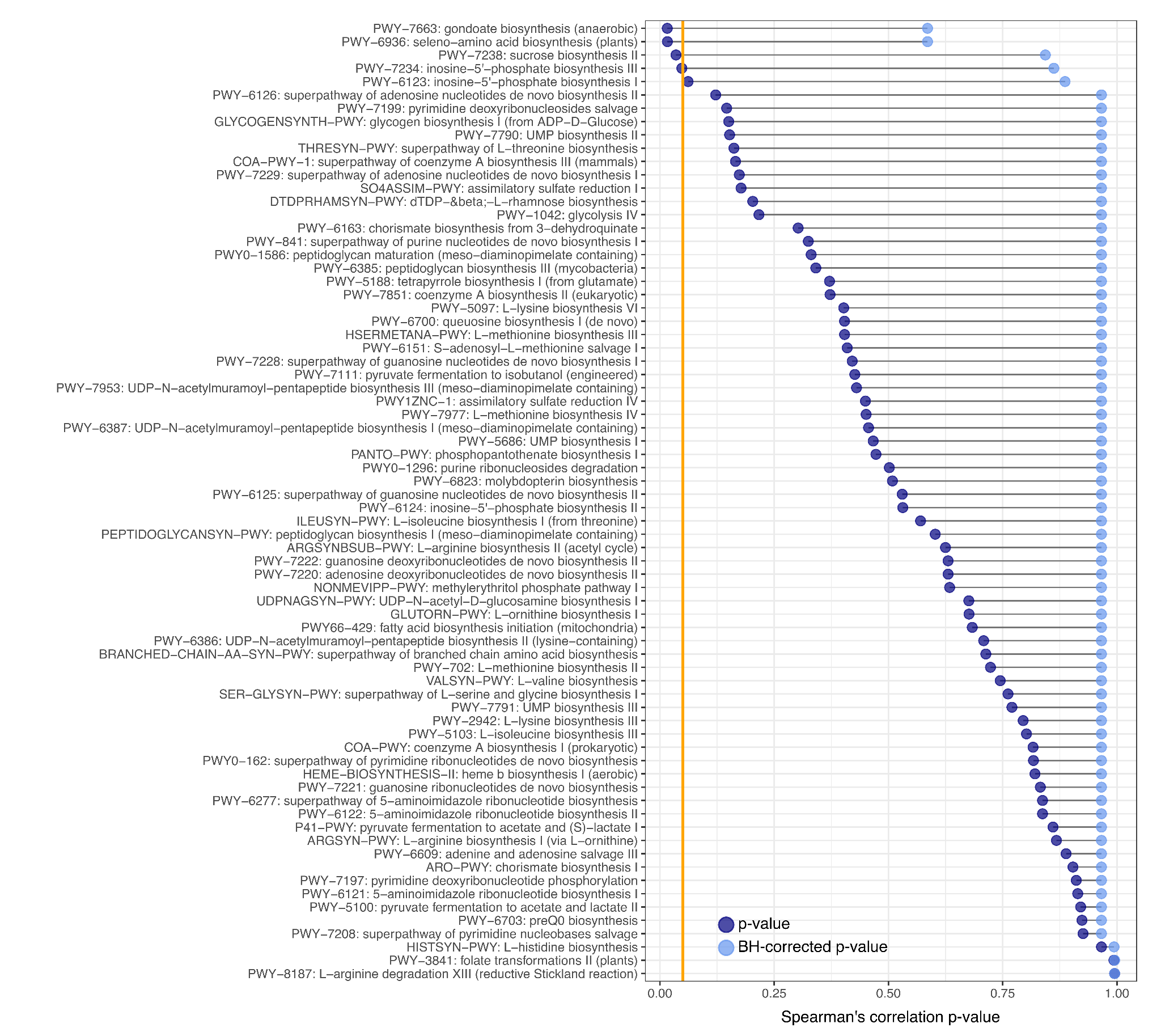

**Extended Data 13**. **Spearman’s correlation p-value for all metabolic pathways identified in both maternal milk and infant stool samples considering all mother infant pairs.** The orange line identifies the significant thresholds (*p*=0.05). P-values were corrected for multiple testing using Benjamin Hochberg correction.

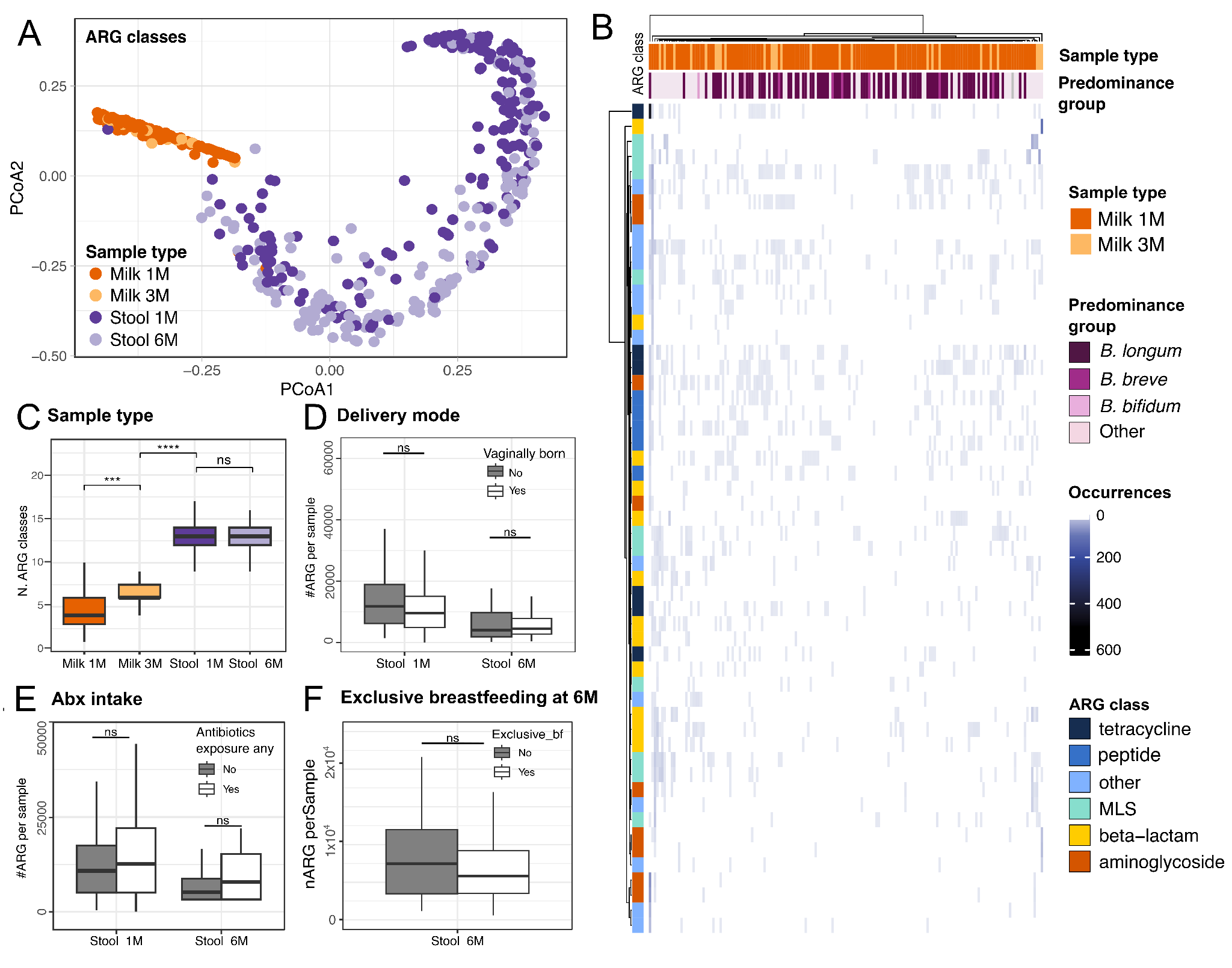

**Extended Data 14**. Antibiotic resistance gene (ARG) profiles across body sites, timepoints, and clinical variables in milk and infant stool samples**.** (A) PCoA of predicted ARGs classes using presence/absence information, as seen by DeepARG. (B) ARGs carriage in milk samples, divided by collection time point, most abundant species (predominance group), number of ARG genes identified and their respective ARGs class. (C) Number of distinct ARG classes identified across body sites and sampling time point. (D-F) Number of ARG genes identified in infant stool samples, divided by delivery mode, history of antibiotic intake and exclusive breastfeeding at 6 months of age, respectively. P-values calculated using t-test, **** for p ≤ 0.0001, ns for non significant.
